# Distinct spatial patterns of neural and physiology-related activity in the human periaqueductal grey during anticipatory social stress

**DOI:** 10.1101/2022.12.29.522243

**Authors:** J.E. Theriault, A. Fischbach, P.A. Kragel, T.D. Wager, L.L. Wald, M. Bianciardi, J.-K. Choi, J. Zhang, K.S. Quigley, L.F. Barrett, A.B. Satpute

**Affiliations:** Department of Psychology, Northeastern University, Boston, MA; Department of Psychology, Emory University, Atlanta, GA; Department of Psychological and Brain Sciences, Dartmouth College, Hanover, NH; Athinoula A. Martinos Center for Biomedical Imaging, Massachusetts General Hospital, Charlestown, MA; Department of Radiology, Massachusetts General Hospital and Harvard Medical School, Charlestown, MA; Department of Surgery, University of California: San Francisco, San Francisco, CA; VA Bedford Healthcare System, Bedford, MA; Department of Psychiatry, Massachusetts General Hospital and Harvard Medical School, Charlestown, MA

## Abstract

Periaqueductal gray (PAG) columns mediate affective experience, physiological regulation, and survival-related behavior; yet, only 7T imaging can resolve these small structures in humans. In a social stress task, participants prepared a speech, and we observed (a) bilateral ventrolateral PAG activity, relative to baseline, and (b) distinct spatial patterns of correlation between PAG activity and physiological response (i.e., cardiac interbeat interval, reparation rate, and tonic electrodermal activity).

---

Functional columns within the periaqueductal gray (PAG) have been identified in human and animal research, but limitations in imaging resolution have hindered studies of their functional involvement in naturalistic human behavior. The PAG is a small midbrain structure (Fig. 1a-d) that plays functional roles in antinociception, blood pressure modulation, migraine pain, vocalization, micturition, sexual behavior, affective experience, and survival-related behaviors (e.g., coordinating a fight-or-flight response)^1^. Of these functions, antinociception, blood pressure modulation, and survival-related behaviors show different (and sometimes opposite) effects following stimulation of dorsolateral and ventrolateral PAG columns^1,2^. Stimulating dorsolateral or lateral PAG elicits a suite of behavioral and physiological reactions that have been interpreted as a defensive reaction^3^—e.g., in rats, it elicits “activity bursts”, such as explosive running and jumping^4,5^, and in cats, it elicits redirection of blood flow from viscera to hindlimb muscles and increases in heart rate, blood pressure, and respiratory rate^1,6^. By contrast, ventrolateral PAG stimulation in rats elicits “freezing” behavior, a species-specific response to threat that is thought to prevent detection when predators are detected^3,7^. In humans, blood pressure is increased and decreased, respectively, by electrical stimulation near dorsolateral and ventrolateral PAG^8^. Human patients also report intensely unpleasant affective sensations after PAG stimulation, which have been hypothesized to result from ascending sensory activation relayed from lateral/ventrolateral PAG^1^. Prior human functional MRI (fMRI) studies at 3T (Tesla) observed BOLD (blood-oxygen level-dependent) signal intensity increases (i.e., *BOLD increases*) in PAG during the anticipation of threat^9–12^; however, 3T studies cannot spatially resolve PAG columns, or even clearly separate the PAG from the cerebral aqueduct it surrounds, and the few existing 7T studies of the human PAG did not detect bilateral columnar effects^13,14^, likely due to their smaller participant sample size. The present work uses a large sample of participants (N = 90), scanned at 7T with simultaneous physiological recording to help resolve fine-scale spatial distinctions in human PAG function during a naturalistic social stress task.

**Figure 1.**
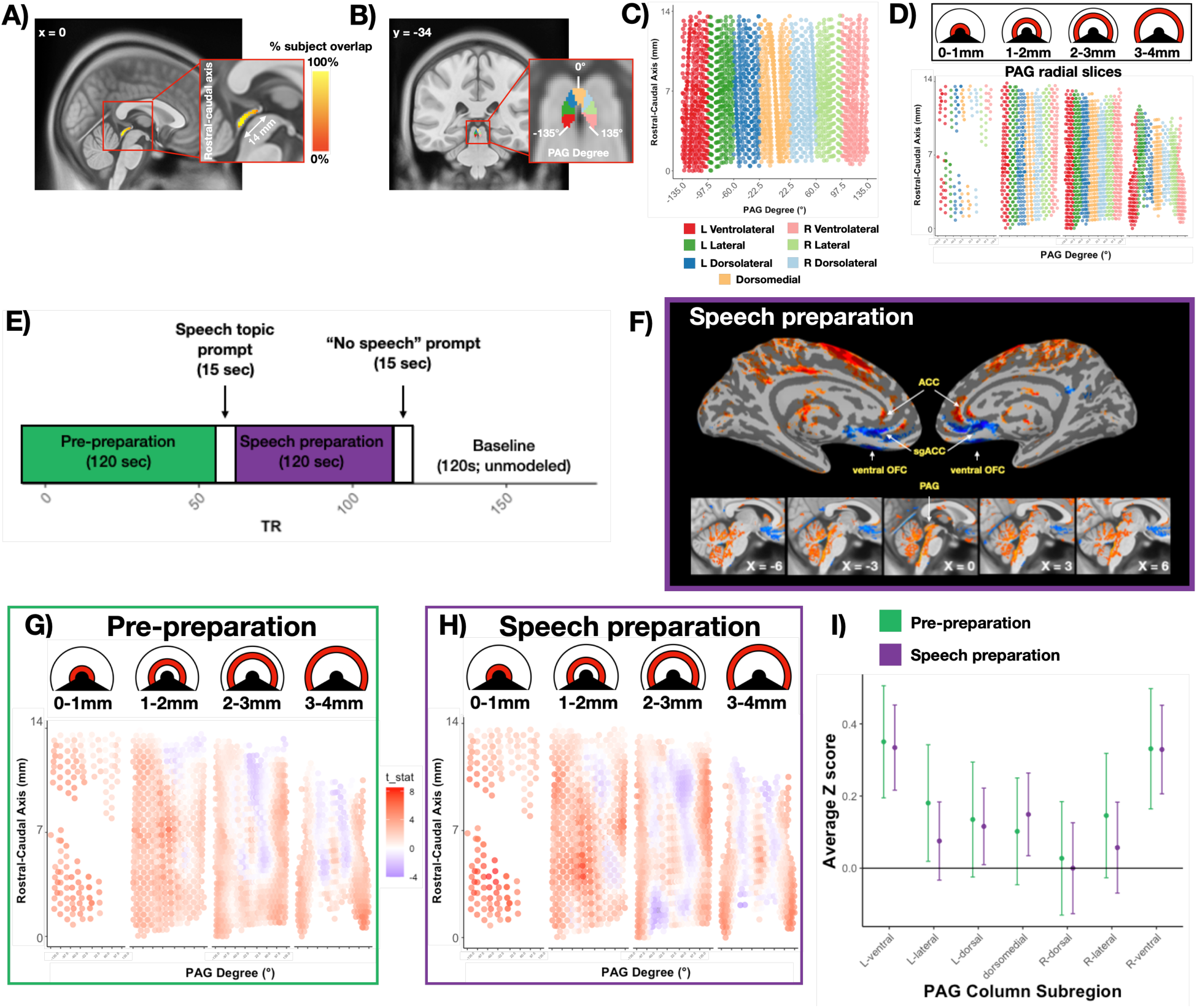
PAG identification, segmentation, and task-elicited BOLD signal intensity. A) The PAG is a small midbrain structure surrounding the cerebral aqueduct. Using GLM residuals, we created subject-specific PAG masks by identifying high variance voxels in the cerebral aqueduct and building a PAG mask around them (see Methods). These subject-specific masks were warped into alignment to substantially increase mean % overlap across subjects (Dice coefficient) from 62.0% [95% CI: 34.3%, 75.1%] to 80.7% [95%CI: 69.3%, 87.3%]. The panel plots % overlap in a sagittal slice (x = 0). B) x/y/z coordinates in group-level PAG mask were geometrically transformed into radial degrees, and columnar PAG ROIs were drawn (ventrolateral = +/− 97.5-135°; lateral = +/− 60-97.5°; dorsolateral = +/− 22.5-60°; dorsomedial = +/− 22.5°). C) PAG voxel ROIs are displayed on a 2d plane, after reslicing from native space (1.1mm x 1.1mm x 1mm) to 0.5mm isotropic. The x-axis displays PAG radial degrees, and the y-axis displays the longitudinal PAG axis (identified by PCA on the native x/y/z PAG coordinates). Colors indicate columnar PAG ROIs. D) The 2d PAG plot is faceted by radial slices of the PAG to allow easy visualization through the entire cylinder—e.g., the 0-1mm column shows voxels abutting the cerebral aqueduct, and the 1-2mm column shows voxels 1-2mm from the aqueduct center. E) Task design. Subjects were told before the scan run that they would be asked to give a seven-minute speech, which would be recorded and evaluated, and for which they would have two minute to prepare after being given the topic. In the pre-preparation period, subjects were aware they would have to give the speech, but did not yet know the topic. In the speech preparation period, participants knew the topic and believed they would speak at its end. Previous studies using this task combined the pre-preparation and final baseline into a single baseline, but we separated them given that the two baseline periods might be experienced as psychologically distinct^15,16^. F) Midline visualization of the whole-brain speech preparation contrast. We observed midline patterns of BOLD intensity consistent with sympathetic activation (top panel), and BOLD signal increases throughout the PAG (bottom panel). Subject-level contrast images were smoothed 1.5 mm and warped to align subject-specific PAG masks (see Methods) before group-level analyses. G) Pre-preparation 2d PAG BOLD signal intensity visualization. PAG voxel z-scores were extracted from each subject’s PAG-aligned, unsmoothed, whole-brain contrast using the group-level PAG mask. After masking, subject PAG voxels were resampled to 0.5mm isotropic and smoothed 1mm. In each voxel, a t-test compared the population of z-score normalized subject estimates to 0. T-tests are plotted using the heatmap, with negative scores in blue and positive in red. Z-score normalized subject estimates were used to correct for heterogeneous variance across PAG voxels, which was especially apparent in voxels bordering the cerebral aqueduct (Fig. S4). H) The same 2d PAG BOLD signal intensity visualization is displayed for the speech preparation period. Note the clear bilateral bands of positive t-tests in the ventrolateral columns. I) PAG column averages and 95% confidence intervals across the subject population, averaging subject ROI z-scores across voxels within each columnar PAG ROI (Fig. 1d).

We used a social stress task, asking participants to prepare for a speech (Fig. 1e), which has elicited PAG BOLD increases in prior work^15,16^, and is a naturalistic manipulation of fear and anxiety-provoking situations that humans regularly encounter. Theoretical models of fear and anxiety in humans have suggested that the PAG is active in response to imminent threats^9–12^, but PAG columns have also been hypothesized to support more specific survival-related responses, with ventrolateral PAG controlling conditioned defenses after a predator is encountered (e.g., “freezing”), and dorsolateral PAG controlling more imminent fight-or-flight reactions, when the predator must be fought or escaped^3,17–20^. Given this, if columnar differences in BOLD increases are observable at 7T during anticipatory social stress, then prior work implies that they would most likely be observed in ventrolateral PAG, given the more distant nature of the social threat. Following prior work^14^, we created subject-specific PAG masks (using the cerebral aqueduct as a landmark; see Online Methods) and aligned these masks to a common space for group-level comparisons.

Consistent with this hypothesis, we did observe BOLD increases in bilateral ventrolateral PAG (Fig 1g-i) as participants prepared to deliver a speech. This result was observed both during the speech preparation period, where participants were told the speech topic (“Why am I a good friend”) and given two minutes to prepare a seven-minute speech (Fig. 1h), and during a prepreparation period, where participants knew they would be asked to give a speech but did not yet know the topic (Fig. 1g). All contrasts were performed in comparison to a baseline period, where participants were informed that they would not have to give a speech and were instructed to relax. The majority of participants reported fully believing the deception (80%; n = 72), and the observed increase in ventrolateral PAG during the speech preparation period was only observed in participants that believed they would give a speech (Fig. S1).

The speech preparation period (but not the pre-preparation period) elicited a midline BOLD pattern consistent with observations in previous publications using the speech preparation task^15,16^, and with a sympathetic physiological response (Fig. 1f; Fig S2)—i.e., BOLD increases in anterior cingulate cortex, and BOLD decreases in subgenual anterior cingulate (or ventromedial prefrontal cortex; vmPFC). Note that prior human-based work, using 3T imaging, reported BOLD increases in vmPFC in response to distant or invisible predators and BOLD increases in PAG for imminent or visible predators^9–12^. In contrast, our two-minute delay in advance of the speech (i.e., anticipated social threat) is slower than the slowest approaching predator used prior work^11^, yet elicited the putative signature of an imminent threat: a BOLD decrease in vmPFC and a BOLD increase in PAG.

Because the whole-brain BOLD contrast for the speech preparation period was consistent with a sympathetic physiological response, we next examined physiological channels— respiratory rate (RR), cardiac interbeat interval (IBI), and tonic electrodermal activity (EDA)— which were simultaneously recorded during scanning. Channels were checked for noise using manual inspection and custom software (see Online Methods), leaving a participant sample of 55 for IBI, 48 for RR, and 68 for EDA. During speech preparation, all physiological measures increased relative to the baseline period (Fig. 2a-c), and all but respiratory rate increased relative to the pre-preparation period (RR, *b* = 0.38, *t*(47.00) = 1.00, *p* > 0.3; IBI, *b* = −61.31, *t*(54.00) = 7.45, *p* < 1e^-9^; EDA, *b* = 0.52, *t*(70.00) = 5.17, *p* < 1e^-5^), suggesting that respiratory rates remained elevated from the beginning of the scan until the speech was revealed as a deception (Fig. 2a). The results also confirms that, on average across participants, the task elicited the intended physiological response.

**Figure 2.**
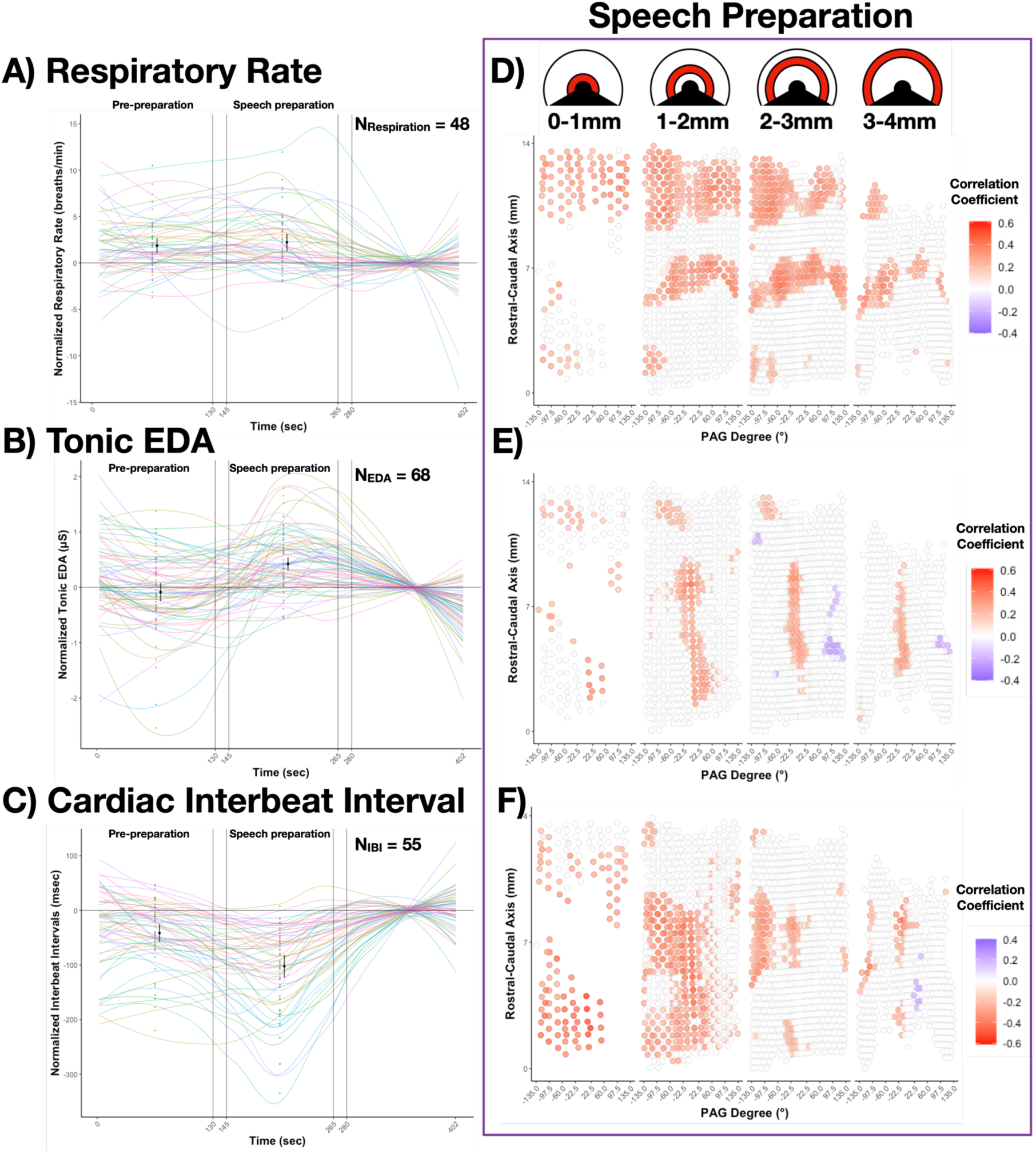
Physiological changes and their across-subject correlation with variance-normalized PAG BOLD signal intensity. A) Colored bands depict loess-function estimates of subject-specific respiratory rates across the task run, normalized to subject-specific average respiration rate in the baseline period. Black dots and confidence intervals indicate subject population means and 95% confidence intervals in pre-preparation and speech preparation periods. The average respiration rate increased in both pre-preparation and speech preparation periods relative to baseline, and did not increase above pre-preparation rates during speech preparation. B) The average tonic EDA activity increased during the speech preparation period relative to baseline, and did not differ from baseline estimates during the pre-preparation period. C) The average cardiac interbeat interval increased relative to baseline in both pre-preparation and speech preparation periods, and increased above pre-preparation rates during speech preparation. D) Across subjects, PAG voxel signal intensity estimates for speech preparation and pre-baseline periods (Fig. S3) were correlated with the average change in respiratory rate from baseline in the same task-period. Correlations were calculated using bootstrap resampling (1000 resamples), and any voxels where the bootstrapped 95% confidence interval included 0 were thresholded. Respiratory rate correlations appeared in clear bands across the rostral and medial segments of the PAG. E) Thresholded cross-subject correlations between PAG signal intensity and tonic EDA change from baseline in the speech preparation period showed a clear band along the dorsomedial column of the PAG. F) Thresholded cross-subject correlations between PAG signal intensity and cardiac interbeat interval change from baseline in the speech preparation period showed a pattern of left lateralization (which was also observed in the pre-preparation period; Fig. S3).

The PAG is a critical relay point between the brain and viscera, and our simultaneous physiological recordings afforded the opportunity to examine *in vivo* physiological correlations with BOLD signal intensity in the PAG. Somewhat to our surprise, we observed clear and distinct spatial patterns of correlation in each physiological channel. Across participants, we correlated PAG BOLD estimates during speech preparation with average change from baseline over the same task period in each physiological channel. Increases in respiratory rate correlated with BOLD in clear bands across the rostral and medial segments of the PAG (Fig. 2d). Increases in tonic skin conductance correlated with BOLD in a clear band along the dorsomedial column (Fig. 2e). Decreases in interbeat interval (i.e., faster heart rates) were correlated with BOLD, largely in left-lateralized medial and caudal segments of the PAG (Fig. 2f). The left-lateralization of IBI was least consistent with the known columnar and rostral–caudal organization of the PAG; but it was also the most internally replicable result of this analysis, as the same topography of correlations was observed in the pre-preparation period (Fig S3). Because both whole-brain task contrasts were left-lateralized as well (consistent with language lateralization during speech preparation; Fig. S2), we speculate that the lateralized IBI-PAG correlation may be driven, in part, by top-down projections from cerebral cortex. By contrast, prior work has suggested that rostral–caudal organization in the PAG stems from somatotopic spinal projections^1,21^, meaning that the rostral–caudal organization of RR-PAG correlations may be more driven by bottom-up spinal projections. These spatial maps of BOLD–physiology correlations should be interpreted with caution: although we defined PAG in relation to subject-specific high-variance aqueduct voxels, some PAG voxels remained more variable than others. To down-weight high variance voxels in group analyses, we used z-score normalized subject-specific BOLD estimates, but despite this, high-variance PAG voxels still generally showed an increase in BOLD-physio correlations, especially for IBI and EDA measurements (but much less so for RR; Fig. S4). This relationship between BOLD–physio correlation strength and voxel variance was likely driven by pulsatile motion in the cerebral aqueduct.

## Acknowledgements

This article was supported by grants from the National Cancer Institute (U01 CA193632 and R01 CA258269-01), the National Science Foundation (BCS 1947972), the National Institute of Mental Health (R01 MH113234, R01 MH109464), the U.S. Army Research Institute for the Behavioral and Social Sciences (W911NF-16-1-019), the National Institutes of Health (NIA-R01AG063982), the Unlikely Collaborators Foundation, and the Roux Institute and the Harold Alfond Foundation. The views, opinions, and/or findings contained in this review are those of the author and shall not be construed as an official Department of the Army position, policy, or decision, unless so designated by other documents, nor do they necessarily reflect the views of the Unlikely Collaborator Foundation.

## Supplemental Materials

**Figure S1.**
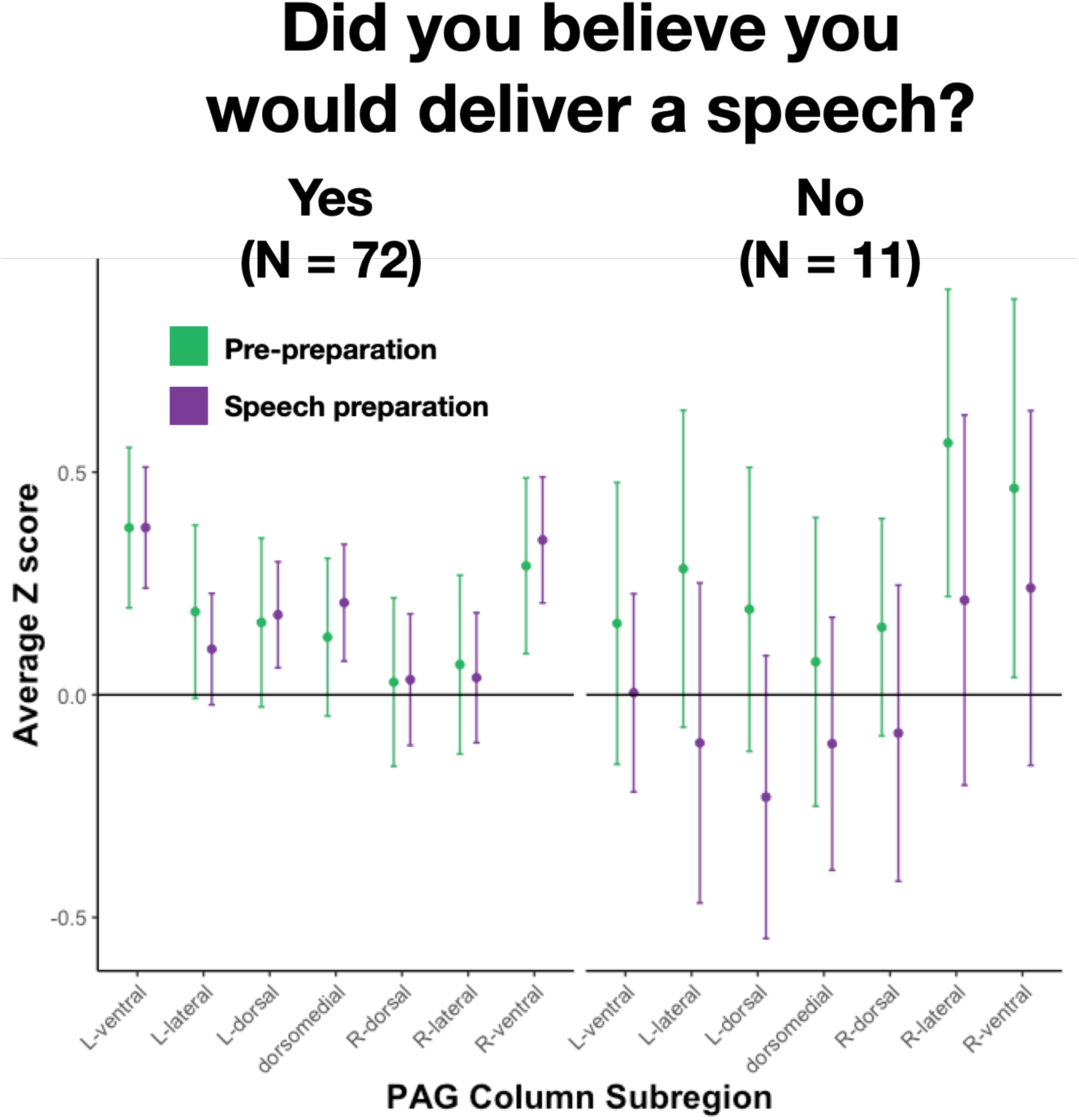
Participants were asked at the end of the study session whether they believed they would deliver a speech in the scanner. Free response answers were recorded, and coded by the experimenter (JT) into clear “Yes” (N = 72) and “No” (N = 11) responses, with any ambiguous responses coded as “Maybe” (N = 6; one participant had no answer recorded). Because of the smaller sample size, PAG column estimates are noisier for subjects where deception failed (i.e., “No”), but there is no clear evidence of a relative increase in ventrolateral PAG, and a clear trend of decreasing PAG signal intensity in the speech preparation period, relative to pre-preparation.

**Figure S2.**
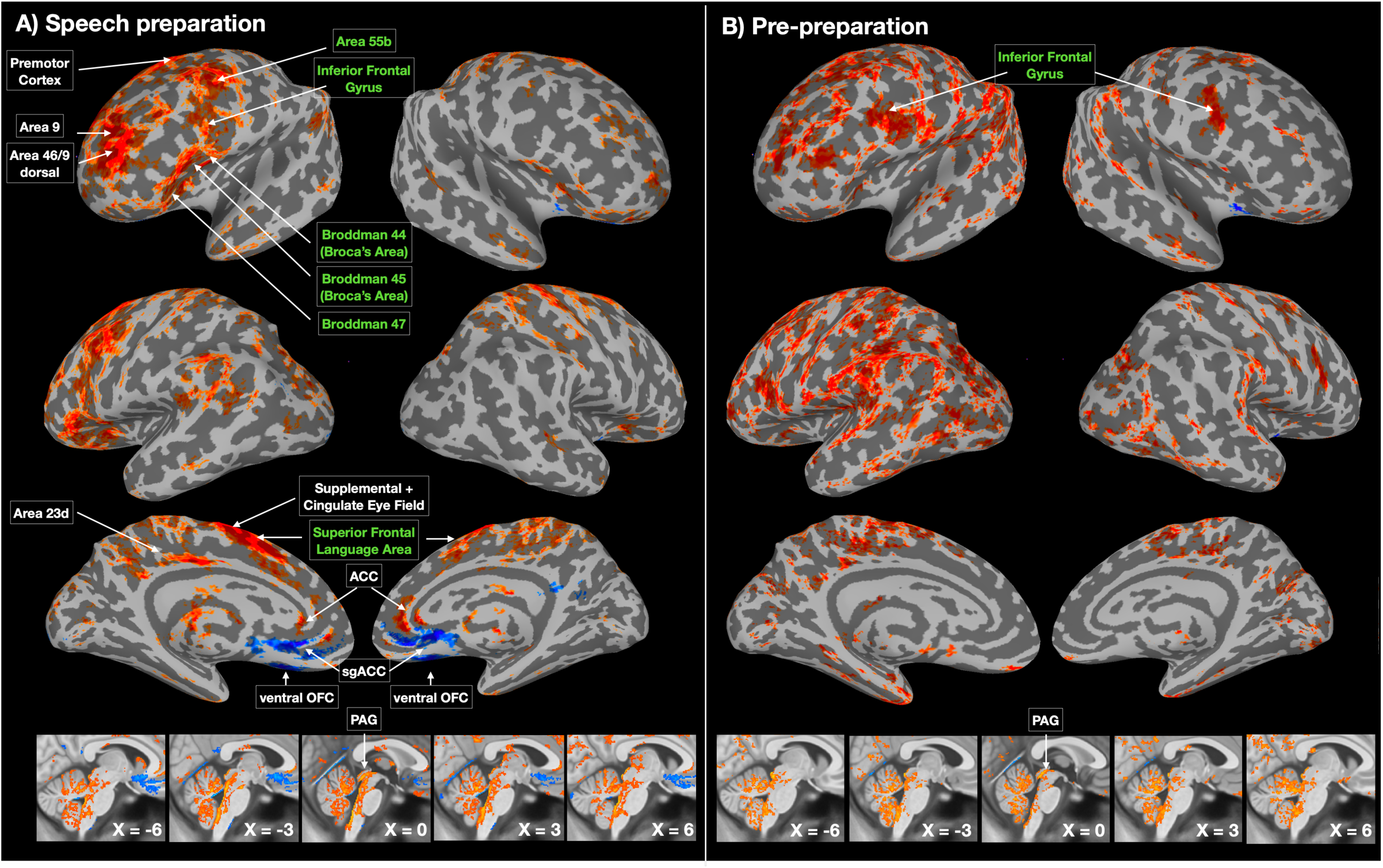
Although midline increases/decreases indicative of a sympathetic response (e.g., increase in ACC, decrease in sgACC) were observed only during speech preparation, left-lateralized cortical changes relative to baseline were observed in both speech preparation and pre-preparation periods. Many clusters observed in the left frontal cortex are associated with language processing (green), which is consistent with the task, given that participants were asked to mentally prepare a speech. Whole-brain estimates were minimally smoothed (1.5mm) and thresholded at a false decision rate of p < 0.05.

**Figure S3.**
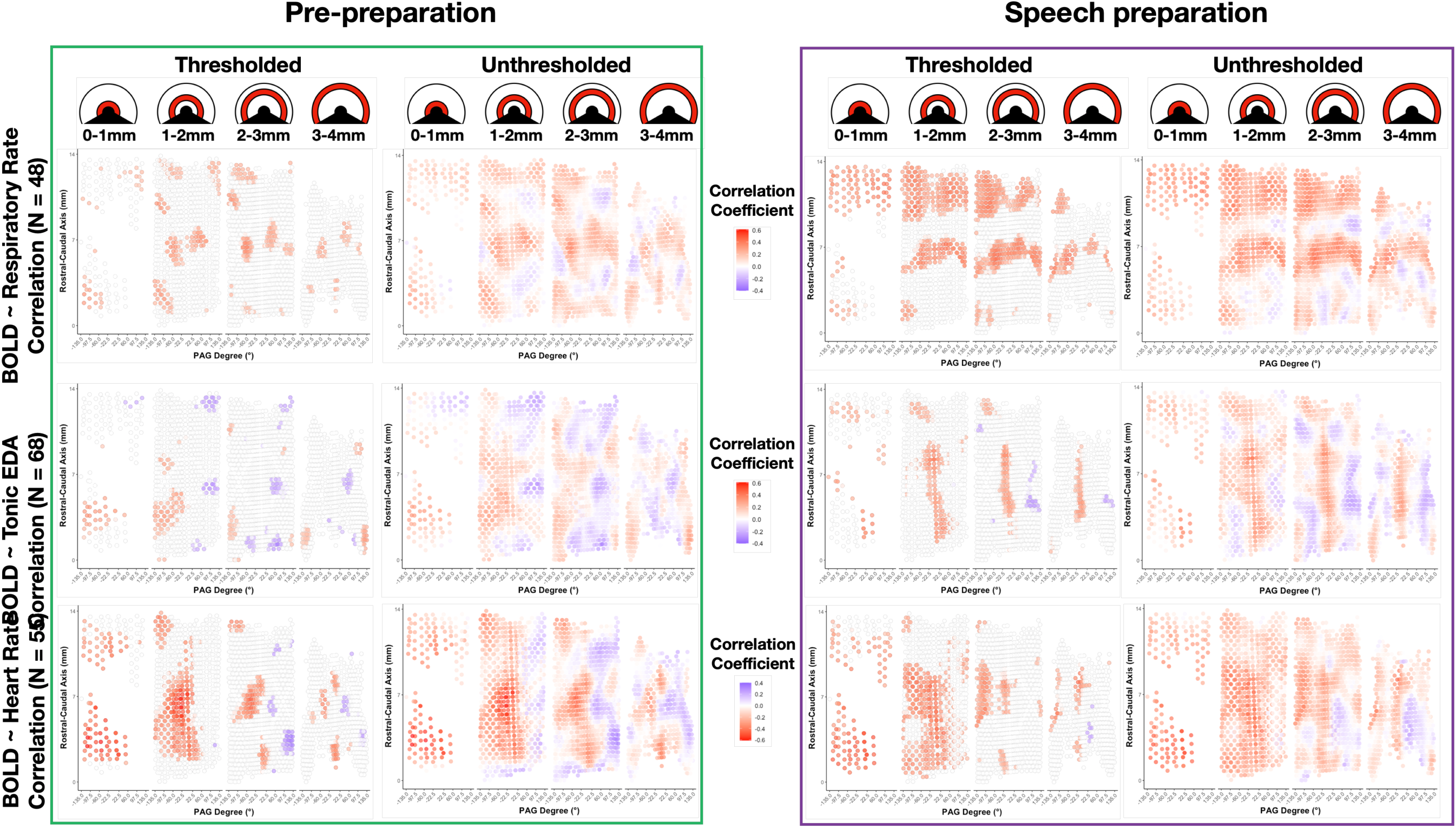
Thresholded and unthresholded PAG-physiology correlation estimates. Correlations were calculated using bootstrap resampling (1000 resamples), and any voxels where the bootstrapped 95% confidence interval included 0 were thresholded. Respiratory rate correlations appeared in clear bands across the rostral and medial segments of the PAG. Note that, for IBI-PAG correlations, the left lateralization replicates between pre-preparation and speech preparation periods (although, given that the effect is stronger in voxels closer to the aqueduct, this effect may also stem from pulsatile motion within the aqueduct; see Fig S4).

**Figure S4.**
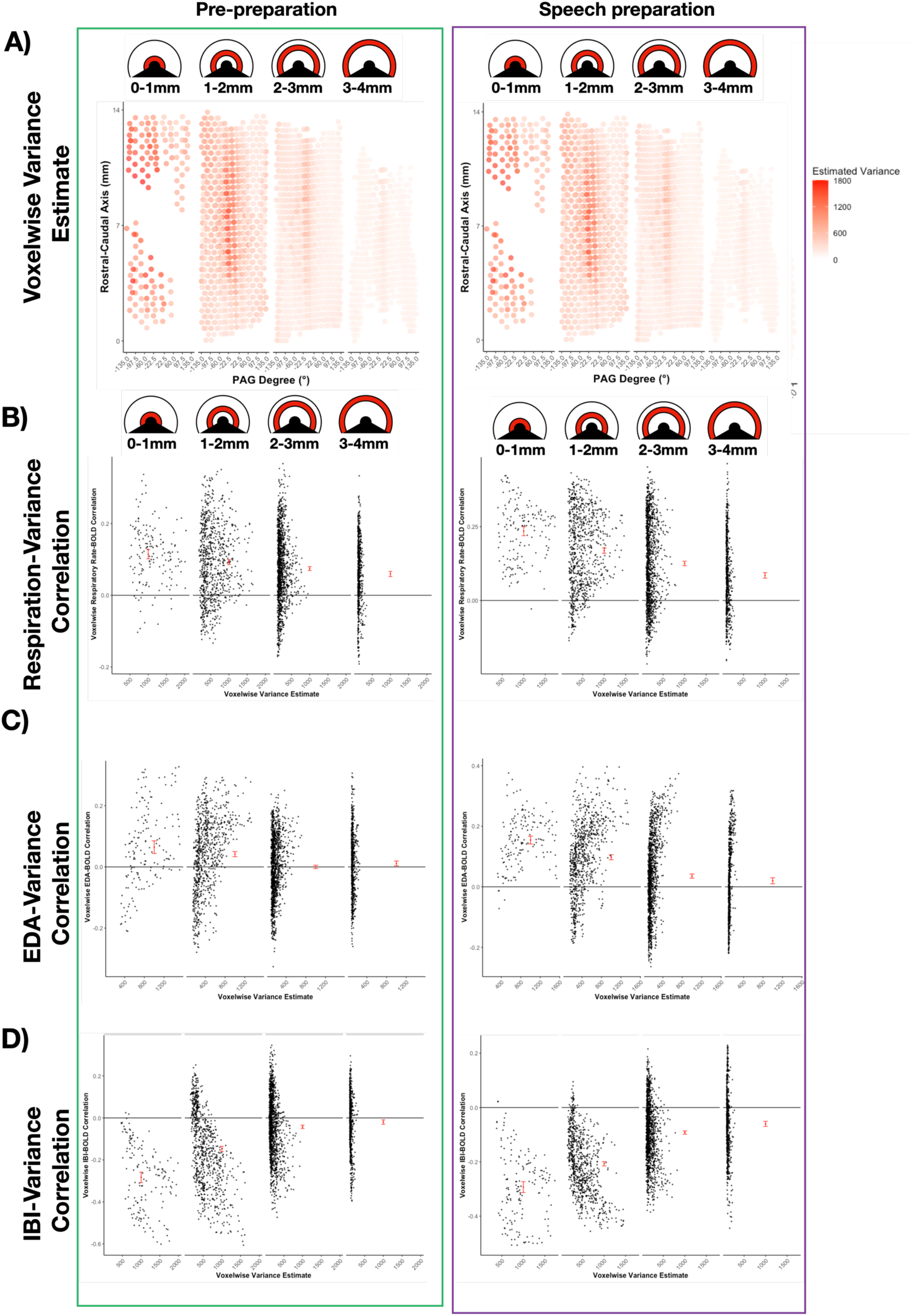
Voxelwise variance estimates and their relationship with BOLD-physiology correlations in PAG voxels. A) Heatmap of estimated variance from the GLM is plotted for each PAG voxel (after resampling to 0.5mm isotropic). X-axis values represent degrees from the PAG midline. Y-axis values represent distance along the rostral-caudal PAG axis. B) Across voxels, group-level correlation estimates between respiratory rate and PAG signal intensity estimates are plotted on the y-axis (see Figs. 2 & S3). The x-axis plots estimated variance for each voxel (i.e., from panel A). Because voxelwise variance estimates clearly differ across radial depth in the PAG, the plot is facetted to plot the variance-correlation relationship in each radial slice. The y-axis mean and 95% confidence interval is displayed in red for each facet. C) The same relationship between voxelwise correlation strength and variance estimates is plotted, with the y-axis now plotting voxelwise across-subject correlations between PAG signal intensity and tonic EDA. D) The same relationship between voxelwise correlation strength and variance estimates is plotted, with the y-axis now plotting voxelwise across-subject correlations between PAG signal intensity and cardiac IBI.

## Online Methods

### Participants

90 participants were included in this study (M_age_ = 26.87 years; SD_age_ = 6.14 years; 38 female, 51 male, 1 non-response). The participant sample was 87.9% non-Hispanic, 61.5% White/Caucasian, 26.4% Asian, and 9.9% Black/African American, and generally well-educated (24.2% had some graduate education, 26.4% had completed college/university, 36.2% had some college/university, 9.8% had completed high school, and 2.2% had not completed high school). Of the 140 participants initially recruited, 12 were excluded due to high motion (see *fMRI preprocessing*), 21 withdrew participation or ended the scan session before completing the speech task, and 16 were excluded due to poor image quality (e.g. subject motion moved large portions of the brain outside the field of view, compromising registration), as assessed by visual inspection in Mango v.4.1 (RRID: SCR_009603). Peripheral physiology recordings were inspected for noise (see *Peripheral physiology recording*) and only clean recordings were included in analysis. Of the 90 participants with usable fMRI data, 55 had clean heart rate data, 48 had clean respiratory data, and 68 had clean electrodermal activity (EDA) data. 26 participants had clean data in all three physiological channels. Participants were recruited from the greater Boston area, were between the ages of 18 and 40 years, were right-handed, had normal or corrected to normal vision, were not pregnant, spoke fluent English, and had no known neurological or psychiatric illnesses. Exclusion criteria included claustrophobia, or the presence of any metal implants. All participants provided written informed consent and study procedures were completed as approved by the Partners’ Healthcare Institutional Review Board.

### Experimental paradigm

Participants prepared to deliver a speech while in the scanner, under the impression that it would be recorded and its merit would be rated by a panel of judges (Cribben et al., 2012, 2013; Lindquist et al., 2007; Lindquist & McKeague, 2009; Wager, van Ast, et al., 2009; Wager, Waugh, et al., 2009). Participants were told that the scan would consist of a 2-minute baseline period where they were to remain still (*pre-baseline*), after which they would have 2 minutes to prepare a 7-minute speech on a presented topic (*speech-prep*), which would be recorded and graded by a panel of experts. In reality, participants did not deliver a speech, and were informed after the speech-prep period that they would not deliver a speech should relax for the remaining 2 minutes of the run (*post-baseline*). The topic (“Why I am a good friend”) was presented for 15 seconds between pre-baseline and speech-prep periods, and the text informing participants they would not give a speech appeared for 15 seconds between speech-prep and post-baseline periods. All instructions were presented using the Psychphysics Toolbox (RRID:SCR_002881, Kleiner et al., 2007) in MATLAB (RRID:SCR_001622, MathWorks).

### fMRI acquisition

Gradient-echo echo-planar imaging BOLD-fMRI was performed on a 7 Tesla MRI scanner at the Athinoula A. Martinos Center for Biomedical Imaging at Massachusetts General Hospital (MGH), Boston, MA. The scanner was built by Magnex Scientific (Oxford, UK), with the MRI console, gradient and gradient drivers, and patient table provided by Siemens. A custom-built 32-channel radiofrequency coil head array was used for reception. Radiofrequency transmission was provided by a 20etonable band-pass birdcage coil. Functional images were acquired using a GRAPPA-EPI sequence (GRAPPA acceleration factor = 3, TE = 28ms, TR = 2.34s, flip angle = 75°, 123 axial slices, A > P phase encoding, partial Fourier in the phase encode direction = 7/8). Structural images were also acquired using a GRAPPA-EPI sequence (GRAPPA acceleration factor = 3, TE = 22 ms, TR = 8.52 s, flip angle = 90°, 126 axial slices, A > P phase encoding,, partial Fourier in the phase encode direction = 6/8). This structural EPI image was reconstructed (via freesurfer and custom scripts) into a T1-like image, which improved anatomical-functional registration and reduced blurring of functional signals by ensuring that anatomical and functional images had similar spatial distortions (Renvall et al., 2016). In both structural and functional images, voxels were 1.1mm isotropic (0mm gap between slices, FOV = 205 x 205mm^2^), echo spacing was 0.81ms, and bandwidth was 1415 Hz per pixel.

### fMRI preprocessing

Preprocessing of the anatomical and functional data was performed using the fmriprep pipeline, version 1.1.2 (Esteban et al., 2019; Esteban, Markiewicz, Goncalves, et al., 2020; RRID: SCR_016216), a Nipype-based tool (Esteban, Markiewicz, Johnson, et al., 2020; Gorgolewski et al., 2011; RRID: SCR_002502). Pipeline details can be found at https://fmriprep.org/en/1.1.2/workflows.html. Each T1w (T1-weighted) volume was corrected for INU (intensity non-uniformity) using `N4BiasFieldCorrection` v2.1.0 (Tustison et al., 2010). Subject brain masks were computed by dilating a binary image of their skull-stripped T1 image by 2 voxels to remove gaps in coverage. Spatial normalization to the 2009c ICBM 152 Nonlinear Asymmetrical template (Fonov et al., 2009; RRID: SCR_008796) was performed through nonlinear registration with the `antsRegistration` tool of ANTs v2.1.0 (Avants et al., 2008; RRID: SCR_004757), using brain-extracted versions of both T1w volume and template. Brain tissue segmentation of cerebrospinal fluid (CSF), white-matter (WM) and gray-matter (GM) was performed on the brain-extracted T1w using `fast` (Zhang et al., 2001) in FSL v5.0.9 (RRID: SCR_002823). Functional data were slice time corrected using `3dTshift` in AFNI v16.2.07 (Cox, 1996; RRID: SCR_005927) and motion corrected using `mcflirt` in FSL (Jenkinson et al., 2002). This was followed by co-registration to the corresponding T1w using boundary-based registration (Greve & Fischl, 2009) with 9 degrees of freedom, using `flirt` in FSL. Motion correcting transformations, BOLD-to-T1w transformation and T1w-to-template (MNI) warp were concatenated and applied in a single step using `antsApplyTransforms` in ANTs using Lanczos interpolation. Physiological noise regressors were extracted using the aCompCor method (Muschelli et al., 2014), taking the top five principle components from subject-specific CSF and WM masks, where the masks were generated by thresholding the WM/CSF masks derived from fast at 99% probability, constraining the CSF mask to the ventricles (using the ALVIN mask; Kempton et al., 2011), and eroding the WM mask using the `binary_erosion` function in SciPy v.1.6.1 (Virtanen et al., 2020). Frame-wise displacement (Power et al., 2014) was calculated for each functional run using the implementation of Nipype. Many internal operations of fmriprep use Nilearn (Abraham et al., 2014; RRID: SCR_001362), principally within the BOLD-processing workflow. For all participants, the quality of brain masks, tissue segmentation, and MNI registration was visually inspected for errors using the html figures provided by the fmriprep pipeline.

### fMRI analysis

To estimate BOLD activity during the speech-prep period, first-level preprocessed functional time series were prewhitened and fit with a general linear model (GLM), using `FILM` (FSL). Four boxcar task regressors were convolved with FSL’s double-gamma hemodynamic response function and included in the GLM to model the onset and duration of the pre-baseline (120 sec.), speech-topic instruction (15 sec.), speech-prep (120 sec.), and no-speech instruction (15 sec.) periods. This modeling strategy differed from previous implementations of this task, which combined pre-/post-baseline periods, but avoids assumptions that participants are equally relaxed in the both periods. Nuisance regressors included framewise displacement, aCompCor components (see *fMRI Preprocessing*), an intercept, and single-TR spike regressors (Satterthwaite et al., 2013), where framewise displacement > 0.5mm. All modeling was performed using wrapper scripts in Nipype v. 1.4.2 (Esteban, Markiewicz, Johnson, et al., 2020). Whole brain contrast maps were masked by the MNI 2009c asymmetric grey matter probability mask (> 20% probability) and thresholded using a false decision rate of p < 0.05.

Long task periods are somewhat unusual for an fMRI design, which typically involves brief events or blocks. On account of this design feature, we omitted translation and rotation motion parameters and high-pass filtering (or equivalently, discrete cosine transform (DCT) functions), instead modeling framewise displacement and a linear trend. Comparisons with alternative models showed that these choices reduced the average variance inflation factor (VIF) across subjects for the speech-prep period (VIF_mean_ = 1.59, VIF_SD_ = 0.13), relative to replacing framewise displacement with translation and rotation (VIF_mean_ = 9. 57, VIF_SD_ = 8.61), or replacing the linear trend with a set of 240 second DCT functions (VIF_mean_ = 7.34, VIF_SD_ = 0.94). VIF remained inflated when pre-/post-baseline periods were combined, as in earlier studies using this task (e.g. Wager, Waugh, et al., 2009).

### PAG identification and alignment

An additional alignment step, beyond the alignment to MNI space, was taken to ensure that BOLD activity estimates in the PAG were aligned across subjects, similar to techniques used in prior work (Kragel et al., 2019; Satpute et al., 2013). The PAG was identified in each subject’s MNI normalized brain and aligned to a common space by: (a) using high-variability model residuals to identify the cerebral aqueduct and create a subject-specific aqueduct mask; (b) drawing a subject-specific PAG mask, dilating the aqueduct mask by 2 voxels (2.2mm), and masking any voxels that were in the original aqueduct mask, that were outside the MNI coordinates range of [-42 < y < −22] and [-14 < z], or that were not in the subject-specific gray-matter segmentation (>50% probability); (c) creating a custom group PAG-aligned template using DARTEL (Ashburner, 2007), which was in approximate MNI space due to first-pass normalization; and (d) warping contrast estimates into the PAG-aligned space. After warping, mean % overlap among subject-specific PAG masks (i.e. the Dice coefficient) increased from 62.0% [95% CI: 34.3%, 75.1%] to 80.7% [95%CI: 69.3%, 87.3%].

The group-aligned PAG template was subdivided into columnar ROIs and ROIs along the PAG rostral-caudal axis (Fig. 1b-d). Voxel x/y/z coordinates were submitted to PCA to identify the PAG longitudinal axis (factor 1), and a geometric transformation on the remaining PCA components gave the radius and degree of each PAG voxel (relative to the y axis, at 0°). PAG columns were defined by their degree range: dorsomedial (+/-22.5°), left/right dorsal (22.5° - 60°), left/right lateral (60° - 97.5°), and left/right ventral (97.5° - 135°), with the ventromedial quarter of the mask excluded from analysis.

### Peripheral physiology recording

All peripheral physiology measures were collected at 1kHz using an AD Instruments Powerlab data acquisition system with MR-compatible sensors and LabChart software. Data were continuously acquired throughout the entire scan session and partitioned for alignment with fMRI data using experimenter annotations in LabChart and scanner TR events. A piezoelectric pulse transducer (AD Instruments) measured heart rate from the left-hand index fingertip. A respiratory belt with a piezo-electric transducer (UFI) measured respiratory rate and was placed around the lower sternum. Wired Ag/AgCl finger electrodes measured electrodermal activity from sensors containing isotonic paste on the left middle and ring fingertips, with signals amplified by an FE116 GSR amplifier.

Physiological time series data were visually inspected for quality in Biopac Student Lab and in custom visualizations using R (v.3.6.2; R Core Team, 2016) and ggplot2 (v.3.2.1; Wickham, 2009). Time series were classified as clean (i.e. clean of artifacts), noisy (i.e. containing artifacts among periods of usable data), and unusable (i.e. containing artifacts and little to no usable data). Contamination of electrodermal activity by respiration was also recorded when present, and a percentage of time course contamination was recorded for each participant and run. Among the 90 participants with usable fMRI data, 55 had clean cardiac data, 17 had noisy cardiac data, 12 had unusable cardiac data, and 7 had no recorded data. For respiratory data, 48 time series were clean, 27 were noisy, 12 were unusable, and 4 had no recorded data. For electrodermal activity, 68 participant time series were clean, 10 were noisy, 9 were unusable, and 4 had no recorded data; and orthogonal to this, 37 time series contained some respiratory contamination (mean % of run contaminated = 59.05%, SD = 32.76%), with 34 of these respiratory-contaminated runs occurring in otherwise clean electrodermal samples. Only clean time series were used in analysis. Among electrodermal activity, clean time series contaminated by respiratory noise were included as this noise was irrelevant to the tonic measures of skin conductance used in analysis.

### Peripheral physiology analysis

Physiological time series data were analyzed using MATLAB toolboxes and custom R scripts. Heart rate was calculated using the PhysIO MATLAB toolbox (Kasper et al., 2017), which used a 0.3 to 9 Hz band-pass filter, and identified cardiac peaks using a two-pass process. The first-pass estimated run average heart rate, detecting peaks exceeding 40% of normalized amplitude, assuming a minimum peak spacing consistent with < 90 beats per minute. First-pass peaks are used to create an averaged template, and the first-pass estimated average heart rate is used as a prior to detect peaks on the second-pass (for more details, see Kasper et al., 2017). PhysIO pipeline peaks were compared with peaks identified in Biopac by trained coders. Discrepancies between the two methods were rare (0.5% of peaks across all runs and participants), and occurred more rarely in the dataset scored by a more experienced coder (0.323% of peaks), compared to the dataset scored by a less experienced coder (0.878% of peaks). Heart rates were smoothed with a 6 second rolling average, downsampled into scan slices, and converted into interbeat intervals. Respiratory rates were calculated using custom R scripts. A 1Hz low-pass filter was applied to respiratory time courses, and local peaks were identified in a sliding 500ms window, removing peaks that fell within 0.5 SD of the run average respiratory value. Respiratory rates were smoothed with a 6 second rolling average and downsampled into scan slices. Tonic electrodermal activity was calculated using Discrete Deconvolution Analysis, as implemented in the Ledalab MATLAB toolbox (Benedek & Kaernbach, 2010).

## Notes

### Competing Interest Statement

The authors have declared no competing interest.

### Summary of Updates

Updated author list.

